# ECLIPSE: Exploring the dark proteome of ESKAPE pathogens through the sequence similarity network of the Protein Universe Atlas

**DOI:** 10.64898/2026.03.30.715302

**Authors:** Surabhi Lata, Dirk W. Heinz

## Abstract

**Motivation:** The accelerating crisis of antimicrobial resistance among the critical, so-called ESKAPE bacterial pathogens demands the urgent identification of novel molecular targets. However, a substantial fraction of ESKAPE proteomes remains functionally uncharacterized, with many genes annotated as encoding hypothetical proteins. These protein sequences often lack significant similarity to known protein families when using conventional homology-based annotation methods and thus remain “dark”. This limits our ability to explore their role in pathogenicity, and it is thus crucial to bridge this substantial gap in pathogen biology by developing novel strategies to illuminate these “dark” regions of the ESKAPE pan-proteomes.

**Results:** We introduce ECLIPSE (ESKAPE Connectome Linkage and Inference for Proteome Sequence Exploration), a network-based computational framework that systematically identifies and prioritises functionally dark protein families in ESKAPE pan-proteomes. ECLIPSE embeds target ESKAPE pathogen proteomes within the global sequence similarity network of the Protein Universe Atlas (Durairaj *et al*. 2023). It detects connected components composed entirely of unannotated proteins, called the “dark proteome”. As a case study, we applied ECLIPSE to a pan-proteome of 3,460,657 protein sequences from 635 strains of *Pseudomonas aeruginosa* (*PA*). ECLIPSE identified 120,985 proteins (4%) residing in completely dark connected components.

Furthermore, we performed a taxonomic diversity analysis using normalized Shannon indices to characterize each dark component by its enrichment in ESKAPE pathogens. The analysis utilized the *evenness (E)* value (see Methods 2.1), which distinguishes *Pseudomonas*-specific (target-specific) from ESKAPE-enriched dark components.

We then developed the Dark Proteome Prioritization Score (DPPS), a composite multi-dimensional scoring framework (see Methods 2.5). It ranks these dark components by biological relevance across four orthogonal axes: (i) functional darkness, (ii) *P. aeruginosa* proportion in the Atlas, (iii) AMR-clade taxonomic restriction, and (iv) conservation across the 635 *P. aeruginosa* strains. This framework outputs a robust four-tier scoring system; the prioritized Tier I components were validated by weight sensitivity analysis and remained stable across 500 Monte Carlo weight perturbations. Structural characterization of one of the top-ranked ESKAPE-enriched dark component revealed that it belongs to the □-barrel fold DUF1302 (PF06980) family for which no experimentally solved three-dimensional structure exists in the PDB. The genomic context analysis indicates that it is co-localized with a LuxR-type transcriptional regulator. Collectively, ECLIPSE identifies evolutionarily conserved, structurally defined, and functionally dark proteins enriched across ESKAPE pathogens; these candidates can further facilitate the experimental characterization of dark proteins as an alternative antimicrobial target.

**Availability and implementation:** The source code and dataset are available for free at:

Github: https://github.com/surabhilata/ECLIPSE.git

Zenodo: DOI: https://doi.org/10.5281/zenodo.21064323

## 1 Introduction

The rise of critical multidrug-resistant ESKAPE pathogens (*Enterococcus faecium, Staphylococcus aureus, Klebsiella pneumoniae, Acinetobacter baumannii, Pseudomonas aeruginosa,* and *Enterobacter spp.*), designated as high-priority by the WHO, poses an immense public health threat (De Oliveira *et al*. 2020, Daruka *et al*. 2025). These bacterial pathogens are the leading cause of nosocomial infections worldwide and are responsible for a disproportionate share of antimicrobial-resistant disease (Miller and Arias 2024). Despite the development of next-generation antibiotics, these organisms continue to “escape” available and novel therapies (Becker *et al* 2006, de Kraker *et al* 2016, Heimann *et al*. 2025). Even with the rapid advancements in genomic sequencing, a significant portion of the ESKAPE pathogen proteome remains “dark” because these proteins often lack detectable sequence homology to known families or are expressed only under specific infection conditions (Ghatak *et al*. 2019). These proteins are frequently annotated as “dark” because of the limitations of automated annotation pipelines and the challenges associated with experimental validation. Despite their unknown functions, they may play important roles in virulence and resistance to the existing therapies. Characterizing this “dark proteome” is therefore essential, as it likely harbors hitherto hidden virulence factors and components of metabolic pathways that could serve as promising targets for the development of alternate antimicrobial strategies.

Previous efforts to characterize unannotated proteins in ESKAPE pathogens have largely relied on high-throughput *in silico* analyses and transposon mutagenesis screens followed by experimental validation (Uddin and Jamil 2018, Geisinger *et al*. 2020, Babic and Kovacic 2021, Gray *et al*. 2024, Abdallah *et al*. 2025). These studies have identified many essential proteins and potential drug targets within the conserved core genome of ESKAPE pathogens. However, a pipeline that systematically characterizes hitherto dark proteins across the ESKAPE pan-proteome remains lacking. By mapping the sequence space covered of over 50 million proteins with high-quality predicted structures in the AlphaFold Database (AFDB) (Varadi *et al*. 2024) as a large complex, sequence similarity network, the Protein Universe Atlas (ATLAS) underlying network enables us to address this annotation gap to look beyond pairwise similarities and to their evolutionary context (Durairaj *et al*. 2023).

Here, we introduce ECLIPSE, a scalable computational framework that embeds ESKAPE pathogen pan-proteomes within the global sequence similarity network derived from the Atlas UniRef50 using MMseqs2 (Steinegger and Söding 2017). It enables the systematic identification and prioritization of evolutionarily isolated (“dark”) protein families across millions of sequences in ESKAPE pathogens. ECLIPSE integrates three analytical layers: (i) Atlas-based darkness classification at the community and component levels; (ii) taxonomic diversity analysis using normalized Shannon indices to characterize the evolutionary distribution of dark components across ESKAPE genera; (iii) and the Dark Proteome Prioritization Score (DPPS), a composite multi-dimensional scoring framework that ranks dark candidates by biological relevance for experimental follow-up.

As a case study, we applied ECLIPSE to 635 strains of *Pseudomonas aeruginosa* (*PA*), a leading cause of opportunistic healthcare-associated infections and one of the most extensively studied Gram-negative bacterial pathogens. Despite decades of research into its gene regulation, quorum sensing, virulence, and antibiotic resistance, almost one-third of the *PA* genome remains functionally dark (Freschi *et al*. 2019, Abram *et al* 2022). We demonstrate that ECLIPSE identifies evolutionarily coherent, structurally defined, and AMR-clade-restricted dark protein families, highlighting tractable candidates for experimental characterization as an alternative antimicrobial target. The code for ECLIPSE is implemented as a modular notebook pipeline, which is freely available. It requires only the adjustment of the input dataset and strain count parameters to be applied to any ESKAPE pathogen.

## 2 Methods

### 2.1 ECLIPSE framework overview

The ECLIPSE (ESKAPE Connectome Linkage and Inference for Proteome Sequence Exploration) framework embeds ESKAPE-specific proteomes into a global protein sequence similarity network derived from the Protein Universe Atlas (Figure 1, Supplementary Pseudocode). Rather than relying solely on pairwise sequence comparisons through methods such as BLASTp or HMM–HMM searches, the framework maps each query protein onto the precomputed Atlas sequence-similarity network (Altschul *et al*. 1990, Söding 2005, Durairaj *et al*. 2023). ECLIPSE integrates multiple orthogonal layers of information into a unified analytical workflow. First, large-scale sequence-similarity mapping using MMseqs2 *easy-search* (Steinegger and Söding 2017) locates each query protein within the Atlas connectome dataset (Durairaj *et al*. 2023), linking it to a UniRef50 community of closely related protein sequences and to its parent connected component. A connected component is a group of proteins that are all linked to each other through chains of sequence similarity and separate from the rest of the network. Within each component, proteins that are most closely connected form smaller groups called communities, thus one component usually contains several communities. Second, connectome-level functional brightness metrics quantify the degree of functional annotation for each community and component, enabling classification of proteins as functionally “dark” or “bright”. Third, we compute a taxonomic entropy metric based on normalized Shannon indices, which we refer to as *evenness* (E), to characterize the evolutionary distribution of dark components, distinguishing lineage-specific, pathogen-enriched, and broadly conserved protein families. Evenness computed as:

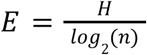

**Figure 1:**
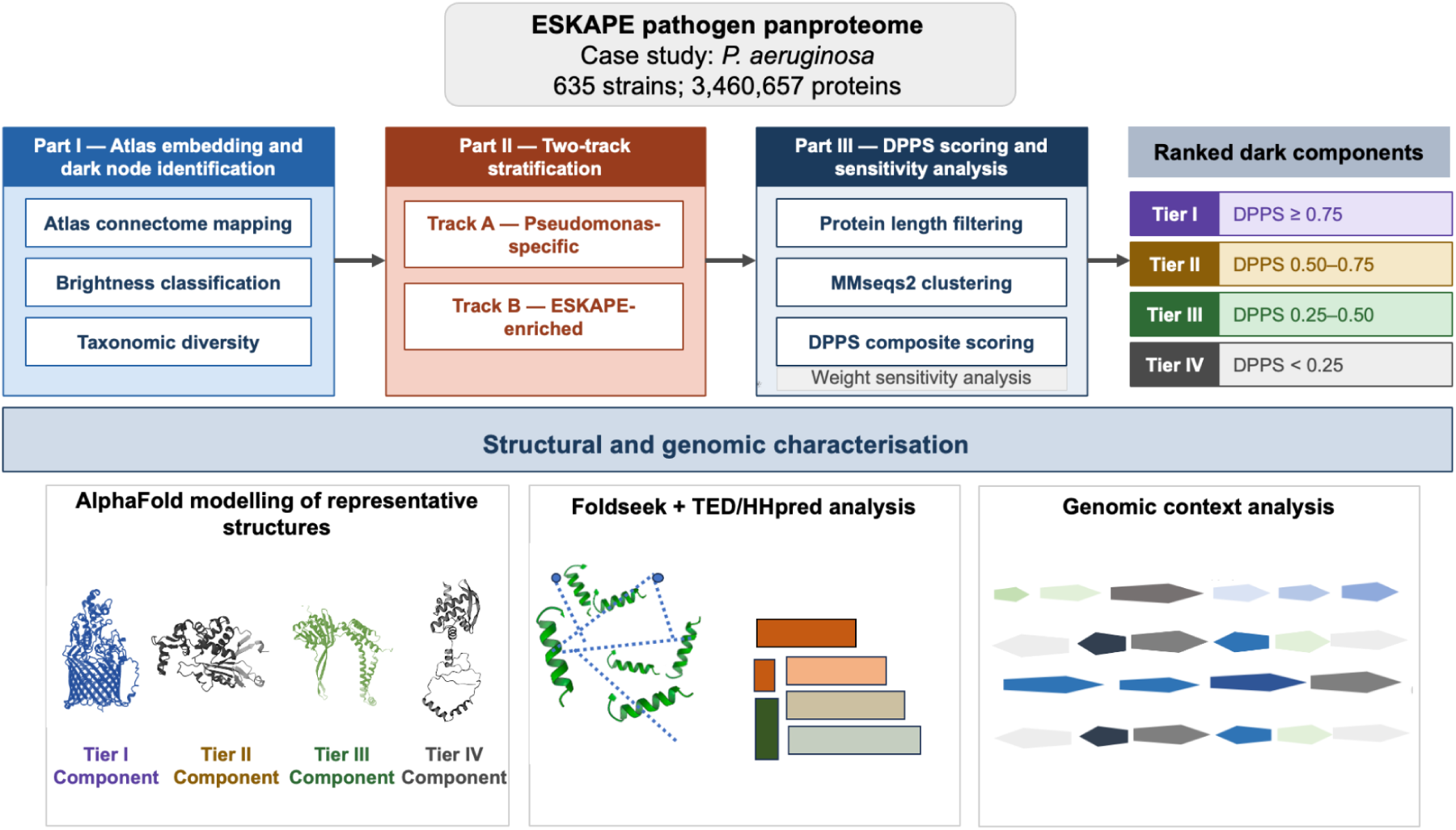
Overview of the ECLIPSE workflow. ECLIPSE accepts protein FASTA files from any ESKAPE pathogen; the *P. aeruginosa* (*PA*) pan-proteome (635 strains, 3,460,657 proteins) is shown as a case study. **Part I** maps each query protein onto the Protein Universe Atlas sequence-similarity network (AFDB90 v.4) using MMseqs2, classifies communities and components as functionally dark or bright by their brightness, and characterises the taxonomic distribution of dark components using a normalised Shannon evenness metric (Methods2.1). **Part II** stratifies dark components into two non-overlapping tracks: Track A (*Pseudomonas*-specific) and Track B (ESKAPE-enriched). **Part III** applies a length filter and MMseqs2 redundancy clustering, then scores each component with the Dark Proteome Prioritisation Score (DPPS) and assesses ranking robustness by a Monte Carlo weight-sensitivity analysis, yielding components ranked into four priority tiers (Supplementary methods). Representative Tier I–IV components are then taken forward for structural and genomic characterisation: AlphaFold modelling of representative structures, Foldseek with TED (The Encyclopedia of Domain) for fold and domain assignment, and genomic-context analysis of conserved gene neighbourhoods. The framework is fully scalable to all ESKAPE pathogens by adjusting the input dataset alone.

where *n* is the number of species and *H* is the Shannon diversity index, computed as

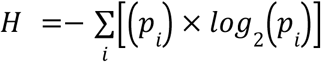

where *i* is the individual species index, and *pᵢ* is the frequency of the protein family found in species *i*.

Following the identification of dark components and their taxonomic characterisation, proteins enriched in ESKAPE pathogens and specifically in *Pseudomonas* were extracted and subjected to the Dark Proteome Prioritization Score (DPPS) framework to rank candidates for downstream structural and functional analyses, which are detailed in the following sections.

### 2.2 Dataset construction

We assembled the ECLIPSE query dataset from the *Pseudomonas* Genome Database by downloading protein FASTA files for all *P. aeruginosa* strains with complete, fully annotated genomes (Winsor *et al*. 2016). Applying this completeness criterion yielded 635 strains, comprising 3,460,657 protein sequences in total. This dataset, used for all subsequent analyses, spans the full spectrum of *P. aeruginosa* genomic diversity relevant to pathogenicity and clinical resistance, including the reference laboratory strain PAO1, the highly virulent clinical strains PA14 and LESB58, and the genetically divergent, multidrug-resistant strain PA7.

### 2.3 Extraction of functional dark nodes

We performed large-scale sequence similarity searches using the mmseqs *easy-search* module of the MMseqs2 version 13-45111+ds-2 (Steinegger and Söding 2017) against the AFDB_90 dataset in the Atlas dataset. It comprises 6,136,321 UniRef50 clusters as of August 2022 (UniRef v.2022_03) and 214,683,829 structural models for most UniProtKB entries available through the AFDB (v.4) network (Durairaj *et al*. 2023). The search was executed with --max-seqs 1, which retains only the best-matching UniRef50 representative per query sequence. Each query is thereby assigned to a single Atlas community and, transitively, to its parent connected component. In the Atlas network, nodes are UniRef50 representatives, and an edge connects two sequences when an MMseqs2 alignment covers at least 50% of the full length of one of the proteins at an *E-value* better than 1×10⁻⁴, with each edge weighted by the *E-value* of the match. Connected components are the maximal sets of nodes mutually reachable through such edges; within each component, communities are densely connected subgraphs identified by asynchronous label propagation (Durairaj *et al*. 2023).

The sequence search result files were mapped to the Atlas community brightness dataset, which contains the brightness percentage of each community per UniProt target ID. Functional brightness is defined as the fraction of full-length close homologues with existing annotations, scored on a continuous scale from 0% (no annotated homologues) to 100% (all homologues annotated). In ECLIPSE, a community is classified as “dark” if its brightness is 0% and “bright” otherwise. A connected component is classified as dark if its median member brightness is 0% and bright otherwise. Brightness values for each community ID were mapped to the corresponding query ID in the search results. The search results were further mapped against the component data file for the AFDB_90 subgraph, and median brightness values were extracted for each query mapped to a component ID. The component dataset summarizes properties of individual connected components, including average brightness, member count, unique protein sequence count, median length, and the number of communities (Durairaj *et al*. 2023).

To benchmark the darkness classification against independent experimental data, three orthogonal ground-truth datasets were assembled for *Pseudomonas aeruginosa (PA)*: (i) virulence (246 genes from VFDB, Victors, and PseudoCAP via the *Pseudomonas* Genome Database v22.1), (ii) antimicrobial resistance (42 genes from CARD v3.2.5 RGI predictions), and (iii) essentiality for viability (75 genes from the GOLD_84 gold-standard essential gene set, cross-validated across five independent Tn-seq screens (Van Maele *et al*. 2026)).

### 2.4 Taxonomic diversity analysis of mapped dark components

To characterize the taxonomic diversity of each dark component, we isolated a subset of proteins from the Atlas data corresponding to ESKAPE species and quantified diversity using three metrics. (i) *ESKAPE relative evenness* estimates taxonomic evenness within each component based on a normalised Shannon index, computed after aggregating all ESKAPE genera (*Acinetobacter*, *Enterococcus*, *Escherichia*, *Klebsiella*, *Pseudomonas*, *Staphylococcus*, *Enterobacter*) into a single “AMR genus” label. A value of 0.0 indicates that the component consists entirely of proteins from either a single non-AMR genus or the aggregated AMR genera; higher values indicate increasing taxonomic heterogeneity beyond the AMR clade. (ii) *ESKAPE genus evenness* reflects diversity exclusively within the AMR-genus subset using a normalised Shannon index. A value of 0.0 indicates that all AMR-associated proteins originate from a single genus; values above 0.0 indicate poly-generic composition across ESKAPE genera. (iii) *ESKAPE proportion* quantifies the fraction of total proteins within the component mapping to the defined AMR genera.

Dark components were partitioned into two non-overlapping tracks for downstream prioritization. (i) The *Pseudomonas*-specific track comprises components where *ESKAPE_proportion* = 1.0 and all Atlas genus annotations belong exclusively to *Pseudomonas*. Genus-level rather than species-level filtering was applied because a substantial fraction of genuine *PA* sequences in UniProt carry the non-canonical label *Pseudomonas* sp.; species-level restriction would therefore introduce systematic false negatives. This *PA* specificity at the strain level was confirmed empirically through the S4 sub-score in the DPPS framework. (ii) The ESKAPE-enriched track comprises components where ESKAPE_proportion ≥ 0.5, after explicitly excluding all *Pseudomonas*-specific components to avoid duplication.

### 2.5 Dark proteome prioritization scoring (DPPS)

The (Dark proteome prioritization scoring) DPPS, systematically prioritizes functionally dark components for experimental follow-up. A component-level median sequence length filter (≥ 300 aa) was applied to both *Pseudomonas*-specific and ESKAPE-enriched tracks to selectively target larger domain architectures that are large enough to support reliable AlphaFold2 modelling and Foldseek structural-similarity searches. This filtering threshold is user-adjustable, allowing it to be modified or omitted to focus on smaller sequence spaces. To empirically justify the 300 aa threshold, we compared it against the length distribution of experimentally supported *P. aeruginosa* PAO1 virulence and resistance determinants retrieved from the *Pseudomonas* Genome Database (PGD v22.1; Winsor *et al*. 2016). The PGD virulence annotation comprises 426 evidence entries spanning 368 unique PAO1 locus tags, with evidence aggregated from VFDB, Victors, and PseudoCAP (a single gene may be supported by more than one source). Antimicrobial resistance determinants were retrieved from the PGD CARD v3.2.5 RGI annotations and comprise 58 unique locus tags. Protein lengths were extracted from the PAO1 reference annotation (5,586 CDS). Of 368 unique virulence loci, 210 (57.1%) are ≥ 300 aa (median 350 aa); of 58 resistance loci, 38 (65.5%) are ≥ 300 aa (median 387 aa) (Figure S1A).

Within each track, sequence redundancy was reduced using *mmseqs easy-cluster*, retaining one representative sequence per cluster for scoring. Clustering parameters were 30% identity and 80% bidirectional coverage for the baseline ≥ 300 aa analysis. The DPPS is computed at the connected-component level as the weighted sum of normalized sub-scores: four sub-scores for the *Pseudomonas*-specific track (Track A) and five for the ESKAPE-enriched track (Track B). S1 quantifies functional darkness of the component; S2 / S2b captures *PA (P.aeruginosa)* representation, either from Atlas taxonomy or directly from query-strain conservation; S3 measures AMR-clade restriction as the inverse of taxonomic evenness across ESKAPE genera globally; S4 reflects empirical conservation across the 635 query *PA* strains independent of database annotation; and S5, applied exclusively in Track B, captures components simultaneously enriched within AMR-associated genera. All sub-scores are normalised to [0, 1] before weighting, and weights within each track sum to 1.0. Components were stratified as Tier I (DPPS ≥ 0.75), Tier II (0.50 ≤ DPPS < 0.75), Tier III (0.25 ≤ DPPS < 0.50), or Tier IV (DPPS < 0.25). Sub-score weights encode an explicit ordinal prior over the biological signals most indicative of pathogen-relevant dark families: AMR-clade restriction (S3), strain conservation (S4), *PA*-specificity (S2b in Track A; S2 in Track B), and ESKAPE-enrichment (S5; Track B) are weighted most heavily because they most directly separate conserved, lineage-restricted families from broadly distributed hypothetical proteins. S1 is assigned the lowest weight (*w*₁ = 0.15) because it evaluates to ≈1.0 for all retained components by construction. Full sub-score definitions, weights, and biological rationale appear in Supplementary Methods and (Table S1).

To ensure transparency over the filtered fraction, DPPS was additionally applied to dark components with median length < 300 aa using identical sub-score definitions, weights, tier boundaries, and Monte Carlo parameters, but with a relaxed MMseqs2 *easy-cluster* coverage of 0.5.

### 2.6 Weight sensitivity analysis

The robustness of DPPS (Dark Proteome Prioritization Score) rankings was tested by a Monte Carlo sensitivity analysis 500 alternative weight vectors were drawn from a symmetric Dirichlet distribution (*α* = 1) over the applicable sub-scores, and the DPPS was recomputed for each draw. The fraction of draws in which a component achieved Tier I (DPPS ≥ 0.75) defines its Tier I stability score; components with stability > 0.80 were considered as prioritized dark components.

### 2.7 Multi-level functional annotation and structural inference

To resolve the biological identity of high-priority candidates, the representative sequence for Component 95203 was modeled with AlphaFold2 (Jumper *et al*. 2021), and the model was used as a query for Foldseek (van Kempen *et al*. 2024) in TM-align mode against the PDB and AFDB (alphafold database). The Encyclopedia of Domains (TED) was used for CATH-based domain assignment (Lau *et al*. 2024; Lang *et al*. 2025). High-tier components and their Foldseek hits were clustered in CLANS (Frickey and Lupas 2004) using all-against-all BLASTp at E ≤ 1×10⁻²⁴ with N-linkage clustering (minimum two sequences). Conserved gene neighborhoods were inferred with GCsnap 2.0 (Pereira 2021).

### 2.8 Comparison with existing annotation approaches and computational requirements

ECLIPSE addresses a methodological gap not jointly covered by existing tools, which fall into three categories relevant to dark proteome analysis: (i) Pairwise sequence-search tools (BLASTp, HHblits, HMMER) return ranked alignment lists rather than pan-proteome scale family graphs. Iterative variants (jackhmmer, HHblits) can recover homologs of a single query, but do not natively produce the component-level family partition or cross-pathogen mapping that ECLIPSE requires, and become computationally prohibitive when scaled to all-vs-all comparison of pan-proteomes against protein-universe references such as the AFDB. (Altschul *et al*. 1990, Remmert *et al*. 2011, Finn *et al*. 2011). (ii) Homology- and signature-based annotation pipelines (e.g., eggNOG-mapper, InterProScan, Prokka) assign function using previously characterised homologues, orthologues, or conserved protein domains. Consequently, they have limited ability to annotate proteins lacking characterised homologues or detectable functional signatures, which constitute the primary focus of ECLIPSE (Zdobnov and Apweiler 2001, Seemann 2014, Cantalapiedra *et al*. 2021). (iii) Pangenome and orthology tools (Roary, PanOCT) classify gene families as core, accessory, or unique within a species pangenome but do not embed them into the broader protein universe, and therefore cannot identify families as dark across the known proteome space (Fouts *et al*. 2012, Page *et al*. 2015).

A direct head-to-head benchmark across these categories is uninformative because each tool produces a different output artefact and no curated benchmark of validated dark targets in *P. aeruginosa* exists. ECLIPSE is complementary to these tools rather than competing with them: annotation pipelines resolve the bright fraction, while ECLIPSE delimits and prioritises the dark fraction.

Computationally, the complete three-notebook pipeline runs end-to-end on a standard laptop (16 GB RAM) in ∼34 minutes with peak resident memory of ∼5.5 GB, without requiring HPC infrastructure.

## 3. Results

### 3.1 Quantifying the “dark proteome” of the *P.aeruginosa* (*PA)* pan-proteome

To establish the scale of functional darkness within the (*PA)* pan-proteome, we mapped 3,460,657 protein sequences from 635 phylogenetically diverse *PA* strains against the global sequence similarity network of the Protein Universe Atlas (AFDB_90 v.4, UniRef v.2022_03). Of the 3,457,506 proteins successfully mapped to Atlas communities, 296,755 (9%) were associated with functionally dark communities carrying 0% brightness, indicating that these proteins have no close homologues with functional annotations anywhere in the known protein universe (Figure 2A). At the more stringent connected-component level, which captures evolutionary relationships across broader sequence space, 120,985 proteins (4% of the 2,870,087 component-mapped sequences) fell into entirely dark connected components with a median component brightness of 0% (Figure 2B). These proteins have no annotated relatives at any evolutionary distance captured within the Atlas network, a considerably more rigorous definition of functional darkness than community-level darkness alone.

**Figure 2:**
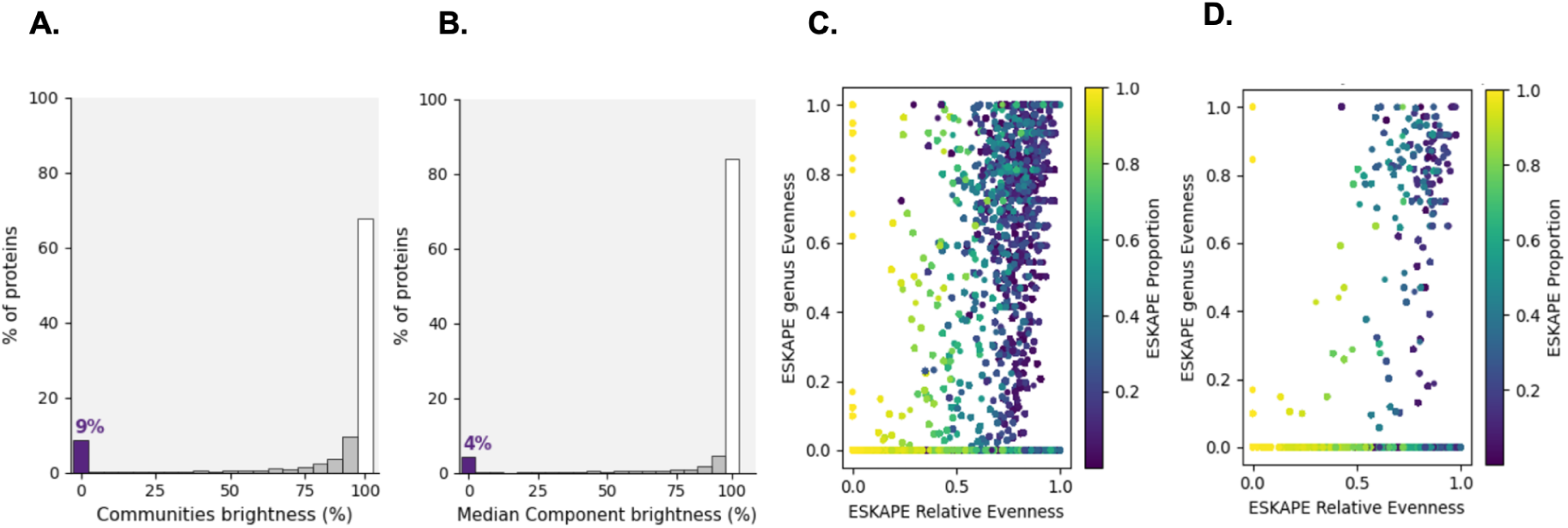
Distribution of functional brightness across communities and components. Histograms illustrate the percentage of (*Pseudomonas aeruginosa*) *PA* pan-proteome sequences mapped to functionally “bright” versus “dark” regions. **A.** Distribution at the level of local protein communities, showing that approximately 9% of sequences reside in 0% brightness (functionally unannotated) regions. **B**. Distribution at the scale of connected components, showing 4% of component-mapped sequences as functionally dark. **C. & D.** Taxonomic diversity and ESKAPE pathogen enrichment of protein components. Scatter plots characterize the evolutionary distribution of mapped components using normalized Shannon indices. **C.** Global view of taxonomic diversity for all components, plotting ESKAPE relative evenness against the proportion of ESKAPE-associated genera. **D.** Targeted analysis of functionally dark components (0% brightness), used to prioritize families that are either strictly Pseudomonas-specific or broadly enriched across the ESKAPE pathogen group for downstream characterization.

As a ground-truth validation, we examined the Atlas brightness classification of experimentally characterised *PA* genes spanning three independent biological axes: virulence, antimicrobial resistance, and essentiality for viability. 99.2% of virulence genes, 100.0% of resistance genes, and 100% of essential genes were correctly classified as residing in bright communities or components. This confirms that ECLIPSE’s darkness classification does not generate systematic false-positive dark calls for genes already known to be functionally important through independent experimental evidence.

### 3.2 Taxonomic diversity of dark components

Scatter plots of ESKAPE genus evenness against ESKAPE relative evenness, coloured by ESKAPE proportion, revealed distinct compositional clusters among all components and within the dark subset (Figure 2C,D). Applying *ESKAPE_proportion* = 1.0 identified 109 ESKAPE-only components while restricting to *Pseudomonas*-only Atlas annotations yielded 83 *Pseudomonas*-specific dark components. Notably, 57 of these 83 carried no Atlas *PA* species annotation, consistent with widespread deposition of genuine *PA* sequences under the label *Pseudomonas* sp. in UniProt rather than biological absence. Applying ESKAPE_proportion ≥ 0.5, after excluding the *Pseudomonas*-specific set, yielded 215 ESKAPE-enriched dark components: protein families with substantial AMR-genus representation across multiple ESKAPE pathogens. These two sets formed the input for all downstream prioritization.

### 3.3 Dark proteome prioritization using the DPPS framework

Applying the DPPS (Dark Proteome Prioritization Score) to both tracks (Figure 3), the ≥ 300 aa filter retained 13 *Pseudomonas*-specific and 61 ESKAPE-enriched components, which clustered to 20 and 102 representative sequences, respectively (Figure 4A). In Track A, two components reached Tier I (Figure 4B). The top component (191318; DPPS = 0.999) is present in 634 of 635 *PA* strains (S4 = 0.998) with perfect AMR-clade restriction (S3 = 1.0) and a combined *PA* evidence score S2b = 0.998, yet zero Atlas *PA* proportion (S2 = 0.000), demonstrating that S2b rescues biologically meaningful candidates suppressed by database annotation gaps (Figure 5). The second Tier I component (194257; DPPS = 0.790) is present in 413 strains with S3 = 1.0. In Track B, five components reached Tier I (mean DPPS = 0.815). Component 92818 ranked highest (DPPS = 0.905), characterized by high ESKAPE enrichment (S5 = 0.901), near-universal *PA* strain coverage across 634 strains (S4 = 0.998), and an Atlas *PA* proportion of 0.722. Component 31745 (DPPS = 0.821) achieved perfect ESKAPE enrichment (S5 = 1.000) and near-universal strain coverage (S4 = 0.998, 634/635 strains). Components 122382 (DPPS = 0.790) and 95203 (DPPS = 0.781) both reached S4 = 0.989 (628 strains) with high ESKAPE enrichment. Component 68574 (DPPS = 0.776) showed the highest Atlas *PA* proportion among Tier I candidates (S2 = 0.840) but more restricted strain coverage (S4 = 0.189, present in 120 strains), suggesting a lineage-restricted distribution within the *PA* collection (Figure 5). A scatter plot of Atlas *PA* proportion against DPPS, coloured by *PA* strain fraction, confirmed that high-ranking components consistently combine Atlas-level *PA* association with broad conservation across the query strain collection (Figure S3). Sub-score heatmaps of all seven Tier I candidates confirmed that top-ranked components score highly across multiple independent axes simultaneously, rather than being elevated by a single dominant sub-score (Figure S4).

**Figure 3:**
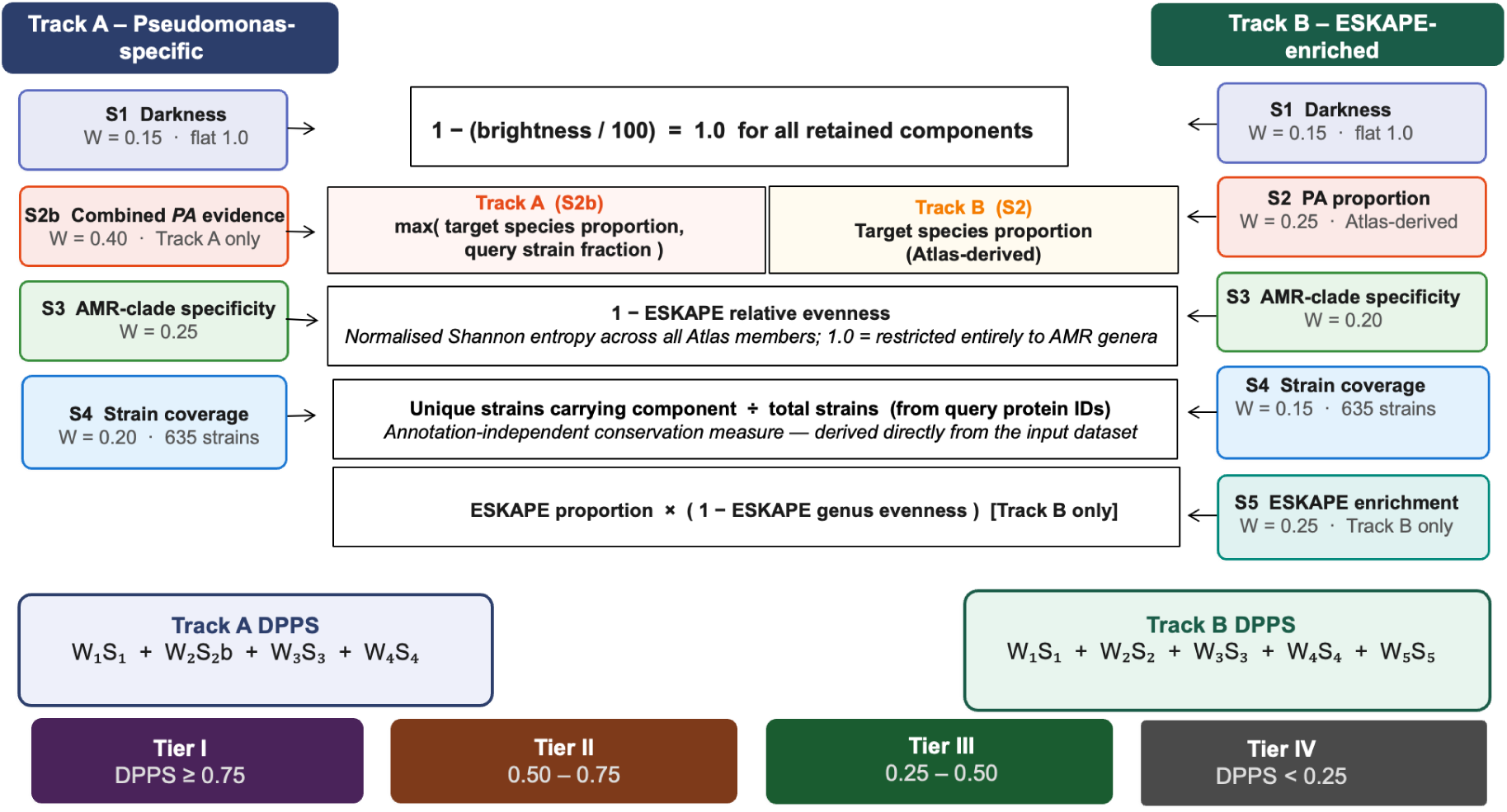
Schematic overview of the Dark Proteome Prioritisation Score (DPPS) framework. DPPS scoring is applied independently to two non-overlapping candidate sets, Track A (*Pseudomonas*-specific, left) and Track B (ESKAPE-enriched, right), reflecting their distinct biological properties and taxonomic composition. Each component receives a weighted sum of sub-scores, all normalized to [0, 1] before weighting. Track A uses four sub-scores (S1–S4) and Track B uses five (S1–S5), with weights summing to 1.0 in both cases. Based on the final score, components are stratified into four priority tiers based on DPPS: Tier I (DPPS ≥ 0.75), Tier II (0.50 ≤ DPPS < 0.75), Tier III (0.25 ≤ DPPS < 0.50), and Tier IV (DPPS < 0.25. The complete mathematical framework is described in detail in the Supplementary Methods and Table S1.

**Figure 4:**
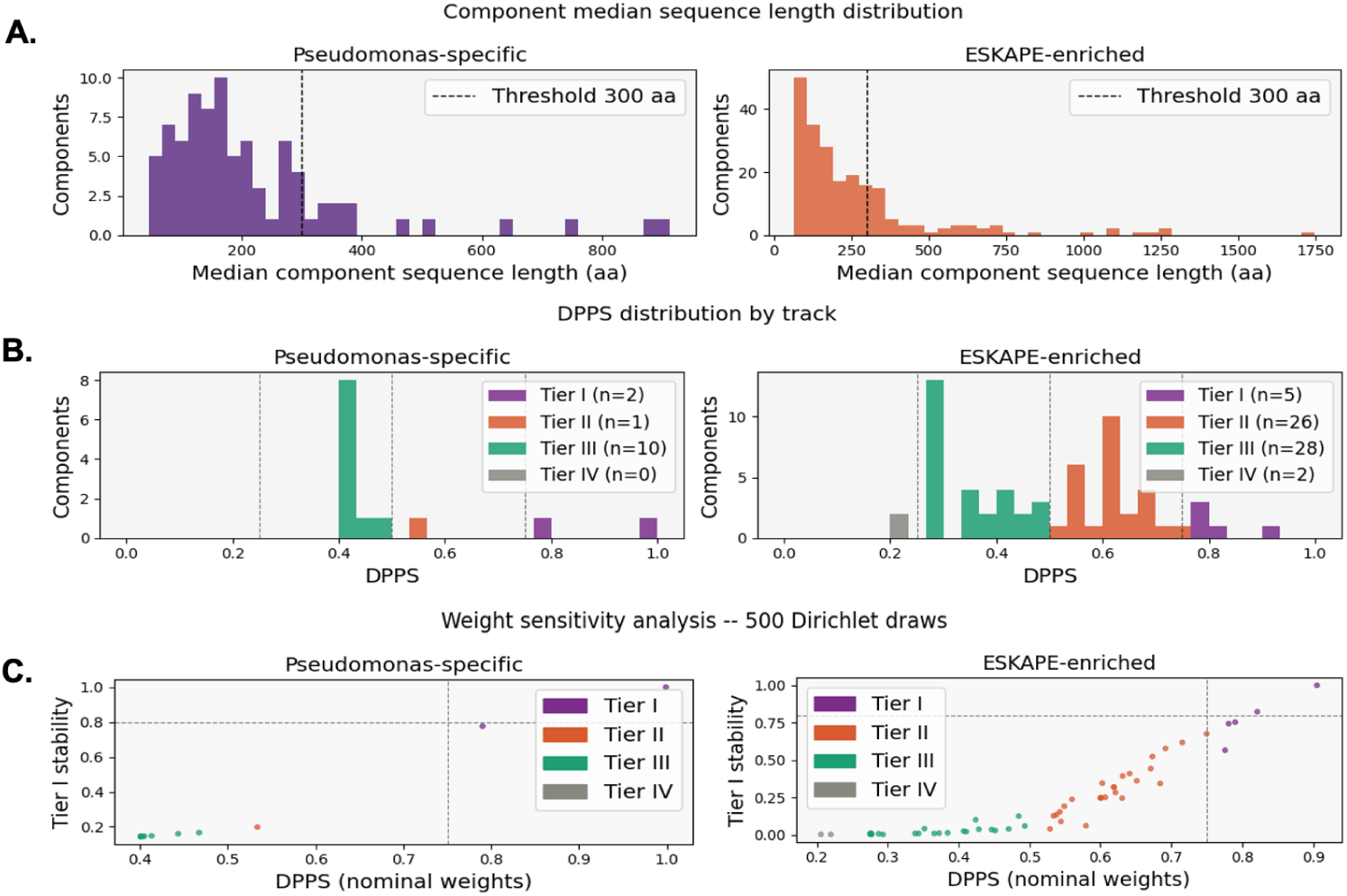
DPPS distribution for *Pseudomonas*-specific and ESKAPE-enriched dark components. **A.** Histograms showing the distribution of median sequence lengths across all dark components in the *Pseudomonas*-specific track (left, purple) and ESKAPE-enriched track (right, orange) before application of the length filter. **B.** Histograms showing the distribution of composite DPPS values for all retained components in the *Pseudomonas*-specific track (left) and ESKAPE-enriched track (right), coloured by tier assignment. **C.** Scatter plots showing, for each component, the nominal DPPS score (x-axis) against its Tier I stability score (y-axis), the fraction of 500 Dirichlet-sampled random weight vectors under which the component achieved Tier I status (DPPS ≥ 0.75). The horizontal dashed line marks the predefined robustness threshold of 0.80; the vertical dashed line marks the Tier I boundary at DPPS = 0.75. Components above both dashed lines are considered robustly prioritized, indicating that their Tier I assignment is independent of specific weight choices.

**Figure 5:**
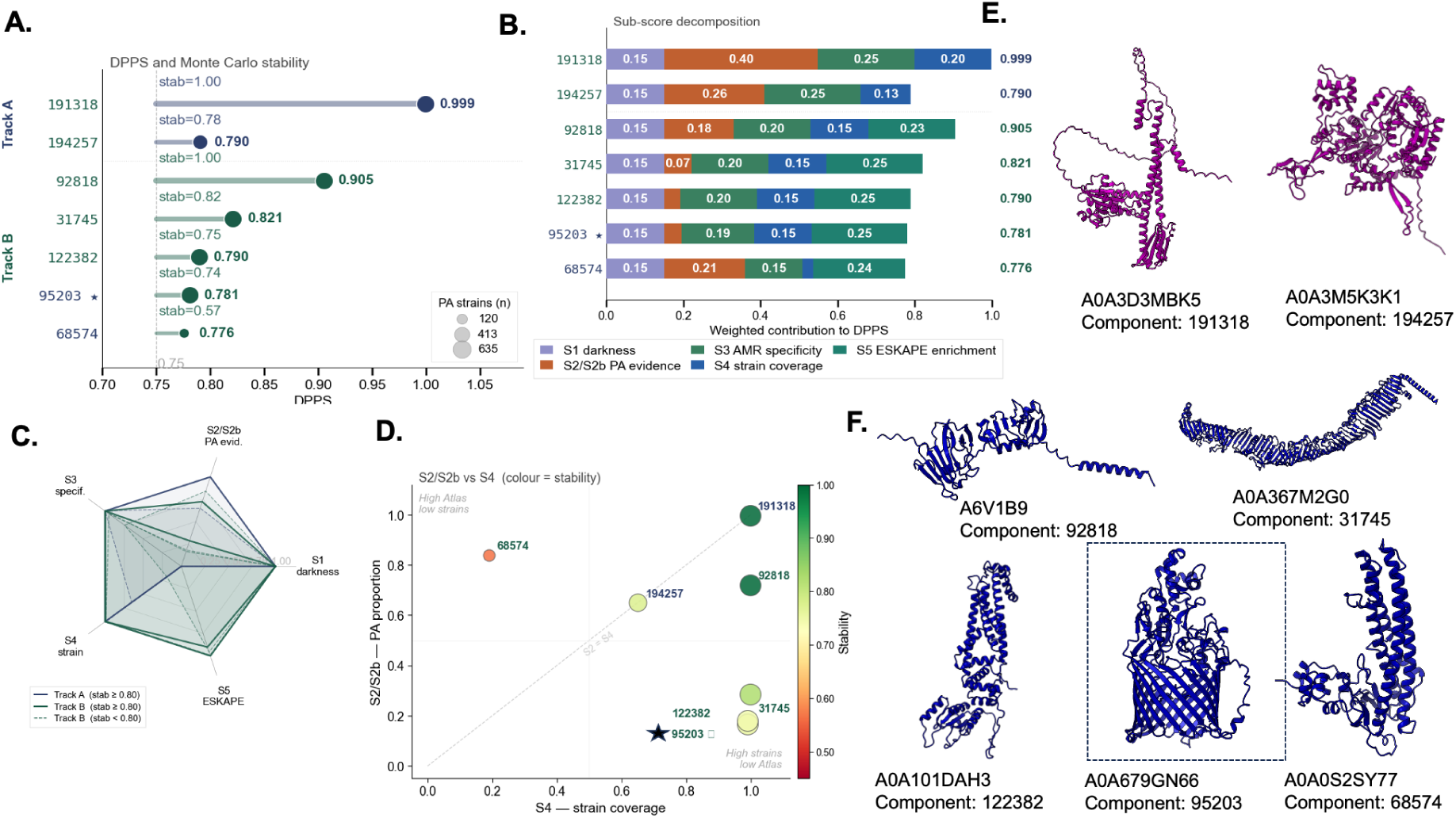
Characterisation of Tier I dark protein candidates from the ECLIPSE DPPS framework. **A.** DPPS lollipop plot: stem position = DPPS value, stem thickness α Monte Carlo stability, circle size α number of *PA* strains carrying the component (120–634 of 635); stability annotated above each stem. Dashed line: Tier I threshold (DPPS = 0.75). **B.** Stacked bar chart showing each sub-score’s weighted contribution to the final DPPS; total bar length = composite DPPS. **(C)** Radar plots of raw (unweighted) sub-score profiles; solid lines indicate a stability ≥ 0.80; while dashed lines indicate a stability < 0.80. **D.** S2/S2b vs S4 bubble plot: bubble area α *PA* strain count; colour represents stability (red < 0.60 → green = 1.00). The diagonal line represents the identity line; points below it indicate prioritization driven by strain conservation rather than Atlas annotation. Component 95203 marked with ★. **E., F.** Representative AF2 (Alphafold2) structures for Track A and Track B Tier I components. The encircled representative is analyzed in Figure 6.

To assess robustness, a Monte Carlo weight sensitivity analysis used 500 Dirichlet-sampled weight vectors. In Track A, component 191318 achieved a Tier I stability score of 1.000, confirming that its top ranking is completely independent of weight choice. Component 194257 reached 0.776, meeting the predefined robustness threshold of 0.80 only marginally, indicating its Tier I assignment is moderately sensitive to weight choice. In Track B, Component 92818 achieved Tier I stability 1.000 and Component 31745 reached 0.824, both exceeding the threshold (Figure 4C). These results demonstrate that the highest-priority DPPS candidates reflect genuine multi-dimensional biological signals rather than artifacts of weight choice, supporting their selection for downstream structural and functional characterization.

To ensure complete coverage of the dark proteome across the full sequence-length distribution, the DPPS pipeline was additionally applied to dark components with median length < 300 aa. In Track A, 70 of 83 components passed this short-protein filter (11,759 proteins, 76 representatives); in Track B, 154 of 215 components were retained (24,374 proteins, 159 representatives). DPPS scoring of the short-protein tracks identified 17 Tier I *Pseudomonas*-specific (mean DPPS = 0.939) and 9 Tier I ESKAPE-enriched components (mean DPPS = 0.809) (Figure S1B–D). AF2 (Alphafold2) models for representative Tier I candidates appear in (Figure S2).

### 3.4 Structural and genomic context characterisation of a top-ranked dark candidate

To demonstrate the biological relevance of DPPS-prioritised candidates, we performed integrated structural and genomic context analysis on Component 95203, the highest-scoring ESKAPE-enriched Tier I candidate by ESKAPE enrichment (S5 = 0.989, DPPS = 0.782), present in 629 of 635 *PA* strains (S4 = 0.991). This component was selected as the primary case study because it combines near-universal conservation across the *PA* pan-proteome, perfect AMR-clade taxonomic restriction (S3 = 1.0), and the highest ESKAPE enrichment score among all Tier I candidates, providing the most compelling multi-dimensional signal for experimental follow-up. The representative sequence of Component 95203 was modeled using AF2 (Alphafold2) and is predicted to form an 18-stranded antiparallel β-barrel with short α-helices capping the outer surface (Figure 6A). Foldseek identified several medium-confidence structural matches in the PDB (probability > 95%, TM-score ∼0.55), with the closest similarities to the *Acinetobacter baumannii* outer membrane carboxylate channel and *AlginateE* from *P. aeruginosa* (Whitney *et al* 2011, Zahn *et al*. 2016). However, differences in the upper cross-sectional region suggest functional divergence (Figure 6A). BLASTp clustering, followed by CLANS analysis, showed that DUF1302 and *AlginateE* homologs form distinct sequence clusters (Figure 6B). Genomic context analysis revealed that DUF1302 genes frequently co-occur with DUF1329, a *LuxR-*type transcriptional regulator, and an *AraC* family transcriptional regulator (Figure 6C). A prior transposon-barcode fitness screen in *Pseudomonas putida* suggested that DUF1302/DUF1329 proteins may participate in fatty acid or ester metabolism and are co-expressed within a LuxR-regulated operon (Thompson *et al*. 2020). Consistently, we observed a conserved association with *LuxR*-type regulators, which are linked to quorum sensing and biofilm formation. AlphaFold–Multimer predicted a DUF1302–DUF1329 complex with a high ipTM score (0.84; Figure 6D).

**Figure 6:**
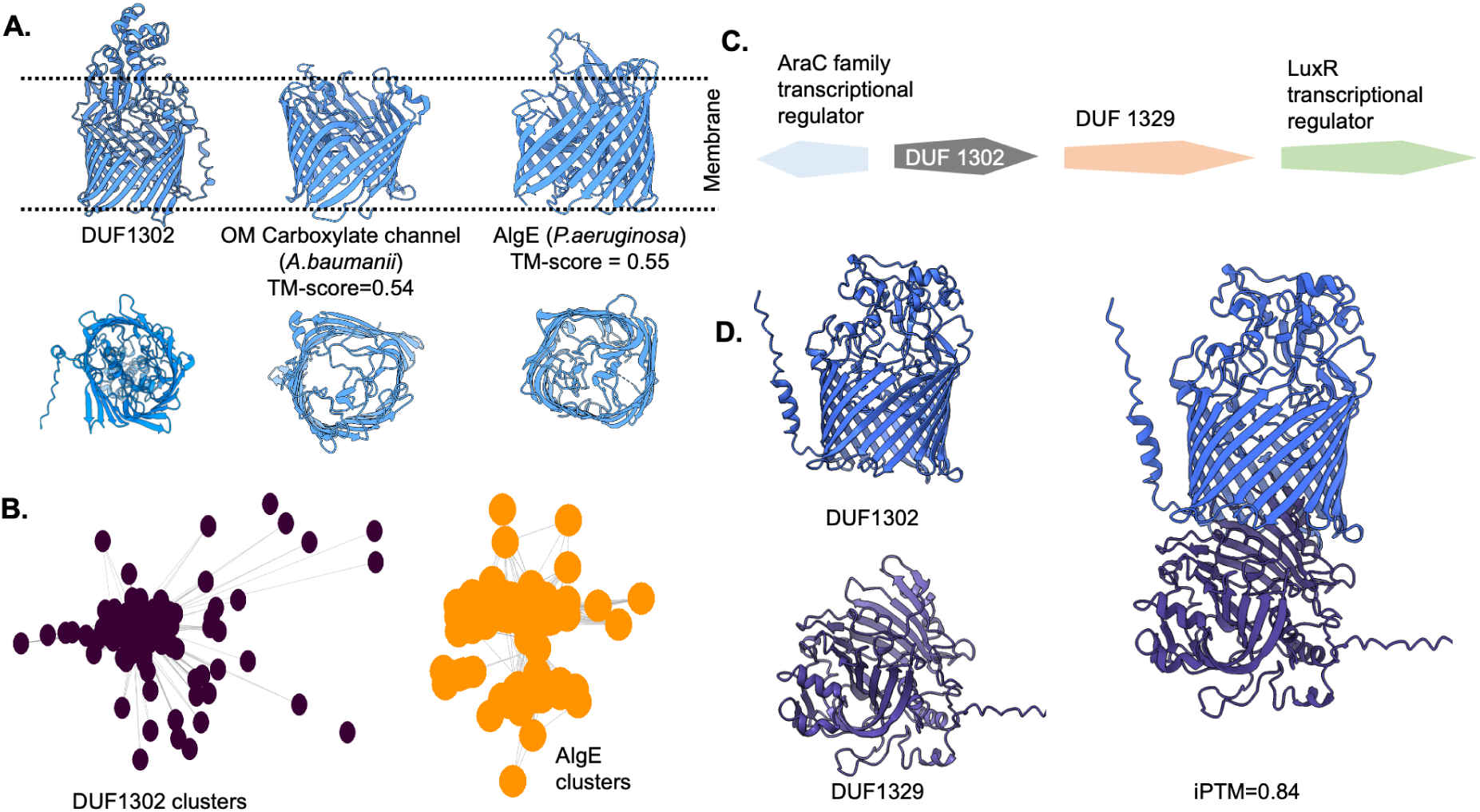
A. Structural and genomic context characterisation of a top-ranked dark component 95203. AlphaFold2-predicted model of the representative sequence for Component 95203 (DUF1302; Pfam PF06980), an 18-stranded antiparallel β-barrel, shown alongside its top Foldseek hits: the *A. baumannii* outer-membrane carboxylate channel (TM-score = 0.54) and *P. aeruginosa* AlgE (TM-score = 0.55). **B.** Sequence similarity network for DUF1302 homologs and Foldseek hits homologs sequences clustered at an e-value of 10^-24^. **C.** Genomic context analysis of flanking genes associated with DUF1302. **D.** Alphafold-multimer complex prediction for DUF1302-DUF1329.

In addition to high-scoring dark proteins, we also analysed the lower-scoring Tier II dark component (Component 71592) (Figure S6).

## 4. Limitations

Limitations of the current study should be noted. The DPPS sub-score weights are biologically motivated but not empirically optimized against a validated training set of confirmed drug targets, as no such curated benchmark exists for dark *PA* proteins. The sensitivity analysis mitigates this limitation by confirming that rankings are robust to weight perturbations, though it does not fully substitute for benchmarking against independent biological datasets such as essentiality screens, virulence-associated genes, or experimentally characterised dark proteins.

We additionally performed an ablation analysis showing that the Tier I candidates are well conserved and that the high-weight sub-scores are the principal contributors to ranking (Figure S5). The central aim of this study is to facilitate the extraction of dark components from the pan-proteomes of ESKAPE pathogens, which can then be further exploited for biological validation.

## 5. Conclusions

The functional characterization of ESKAPE pan-proteomes remains fundamentally incomplete, with a substantial fraction of proteins in even the most intensively studied pathogens carrying no experimentally validated annotation. Here, we present ECLIPSE, a scalable network-based framework that addresses this challenge by embedding ESKAPE pan-proteomes within the global sequence similarity topology of the Protein Universe Atlas. It enables systematic identification of protein families that are dark across the entire known protein universe. Traditional sequence-based pairwise methods such as BLASTp and HMM-based methods, are not scalable to pan-proteomes of millions of sequences against large databases such as the AFDB (Alphafold Database), our network-based approach addresses this gap (Altschul *et al*. 1990), (Durairaj *et al*. 2023). Applied to the *PA (P.aeruginosa)* pan-proteome, ECLIPSE revealed that 4% of component-mapped proteins reside in completely dark connected components. These represent protein families with no characterised relatives at any evolutionary distance captured by the Atlas network, a finding that underscores the scale of unexplored biology in one of the most clinically critical AMR pathogens.

The taxonomic diversity framework introduced here, based on normalized Shannon indices computed across Atlas component membership, provides a quantitative approach for stratifying dark protein families by their evolutionary distribution across the AMR clade. By distinguishing *Pseudomonas*-specific from ESKAPE-enriched dark components, ECLIPSE focuses experimental attention on protein families most likely to perform pathogen-specific functions rather than broadly conserved housekeeping roles. The DPPS (Dark Proteome Prioritization Score) scoring framework further refines this prioritization by integrating four orthogonal biological signals into a continuous ranked list with empirically validated robustness. It addresses a key limitation of binary filter-based approaches that treat all dark proteins equivalently regardless of their conservation, taxonomic restriction, or pathogen association. The structural characterization of Component 95203, a member of the defined Pfam family DUF1302 (PF06980), revealed a β-barrel fold with medium-confidence similarity to known outer membrane proteins. It includes the *A. baumannii* outer membrane carboxylate channel and *AlgE* from *P. aeruginosa* (*PA*) (TM-scores 0.54-0.55). Despite belonging to a defined Pfam family, no experimentally solved three-dimensional structure exists for any DUF1302 member in the PDB, and no direct experimental functional characterization has been performed for this protein family in *PA* specifically. The near-universal conservation of this candidate across 629 of 635 phylogenetically diverse *PA* strains, its enrichment across ESKAPE pathogens, and its conserved genomic co-localization with LuxR-type transcriptional regulators of quorum sensing and biofilm formation collectively suggest specific biological roles in *PA* pathogenicity.

ECLIPSE is designed as a pathogen-agnostic, modular pipeline for ESKAPE pathogens. The notebook implementation accepts the pan-proteome of any ESKAPE pathogen as input, requiring only the adjustment of strain count parameters and input file paths to enable analysis of different species. Applying ECLIPSE across multiple ESKAPE pathogens offers the opportunity to identify conserved dark protein families shared among WHO critical priority pathogens. To promote systematic exploration of ESKAPE dark proteomes, the complete ECLIPSE pipeline, including all analysis notebooks and scoring framework, is freely available for application across the full range of clinically relevant ESKAPE pathogens.

## Acknowledgements

We are grateful to Dr. Joana Pereira (KU Leuven) and all coworkers of Protein Universe Atlas whose published work has inspired us to implement the concept on ESKAPE pathogens. We are grateful to Diana Rapota (Swiss Institute of Bioinformatics, University of Basel) for her help in preparation of jupyter notebook part I and II. We thank Prof. Susanne Engelmann, Prof. Susanne Häussler, Dr. Nicolas Oswaldo Trinler and Dr. Matthias Preusse (all HZI) for valuable discussions.

## Contributions

SL developed the scientific hypothesis, carried out the analysis, and prepared and revised the manuscript draft. DWH established the project’s scope and scale, edited and revised the manuscript.

