## Supplementary material for "ECLIPSE: Exploring the dark proteome of ESKAPE pathogens through the sequence similarity network of the Protein Universe Atlas": SI data

### Supplementary Information

#### Methods

DPPS Scoring:

Scoring was performed at the connected component level, with each component receiving a composite DPPS calculated as the weighted sum of normalised sub-scores. Four sub-scores were used for the *Pseudomonas*-specific track (Track A) and five for the ESKAPE-enriched track (Track B), as described below. All sub-scores are normalised to [0, 1] prior to weighting, and weights within each track sum to 1.0.

S1 (darkness; $w_1$ = 0.15 for both tracks) quantifies functional darkness as 1 − (component brightness / 100), where component brightness is the median Atlas brightness across all UniRef50 members of the component. S1 evaluates to 1.0 for all retained components since only those with a component-level median brightness of 0% pass the upstream filtering step. S1 is included at a modest weight to maintain score normalisation compatibility with relaxed brightness thresholds in future applications and to preserve additive comparability across tracks.

S2b (combined *PA* evidence; $w_2$ = 0.40, Track A only) is defined as max(target species proportion, query strain fraction), where target species proportion is the fraction of Atlas UniRef50 members of the component annotated as *P. aeruginosa* in the Atlas taxonomy files, and query strain fraction is the fraction of the 635 query *PA* strains contributing at least one sequence to the component (equivalent to S4, described below). The max operator was introduced to correct for a systematic annotation artefact specific to the *Pseudomonas*-specific track: inspection of Track A components revealed that 57 of 83 components passing Part II filters showed a target species proportion of zero in the Atlas despite near-universal conservation across the query strain collection, attributable to widespread non-canonical annotation of genuine *PA* sequences as *Pseudomonas* sp. in UniProt rather than as the named species. Without correction, the Atlas-derived species proportion alone would severely penalise highly conserved, genuinely *PA*-specific components. S2b ensures that components are not penalised for annotation gaps by taking the more informative of the two available *PA* evidence signals. S2b receives the highest weight in Track A ($w_2$ = 0.40) because, for components already restricted to the *Pseudomonas* genus by Track A selection criteria, the degree of *PA* specificity within that genus whether measured through the Atlas or directly through query strain conservation  is the most discriminating biological signal for experimental prioritisation.

S2 (PA proportion; $w_2$ = 0.25, Track B only) is the fraction of Atlas UniRef50 members of the component annotated as *P. aeruginosa* in the Atlas taxonomy files, without the max-correction applied in Track A. S2b is not applied in Track B because components in this track are not restricted to a single genus and the annotation gap that motivates S2b is specific to the *Pseudomonas*-centric composition of Track A. S2 weight is reduced relative to Track A ($w_2$ = 0.25) because components in the ESKAPE-enriched track span multiple AMR genera, reducing the discriminating power of species-level Atlas annotation alone; the complementary ESKAPE enrichment signal is instead captured by S5.

S3 (AMR-clade specificity; $w_3$ = 0.25 in Track A, $w_3$ = 0.20 in Track B) is defined as 1 − ESKAPE_relative_evenness, where ESKAPE_relative_evenness is the normalised Shannon entropy computed across all Atlas members of the component after collapsing the seven ESKAPE genera into a single AMR-genus label. A value of 1.0 indicates that all Atlas members of the component belong exclusively to AMR-associated genera across the entire protein universe, while lower values indicate broader phylogenetic distribution. S3 receives the second highest weight in both tracks because AMR-clade restriction globally is the defining biological property distinguishing pathogen-relevant dark proteins from broadly distributed hypothetical proteins.

S4 (PA strain coverage; $w_4$ = 0.20 in Track A, $w_4$ = 0.15 in Track B) is the fraction of the 635 query *PA* strains that contribute at least one protein sequence to the component, computed directly from query protein identifiers by extracting the strain prefix and counting unique strains per component. This metric is threshold-independent and annotation-independent, measuring conservation across the *PA* panproteome empirically from the input dataset rather than from Atlas metadata. Strain identifiers composed of a single token without an underscore separator  indicating a non-standard identifier format incompatible with strain prefix extraction  were excluded from strain counting with a logged warning; four such identifiers were identified and excluded. S4 is weighted at 0.20 in Track A because direct empirical conservation across the query dataset is a strong and annotation-independent confirmation of biological relevance. Its weight is reduced in Track B ($w_4$ = 0.15) because ESKAPE-enriched components span multiple AMR pathogens and *PA* strain-level conservation is one signal among several rather than the primary discriminating criterion.

S5 (ESKAPE enrichment; $w_5$ = 0.25, Track B only) is defined as ESKAPE_proportion × (1 − ESKAPE_genus_evenness), where ESKAPE_proportion is the fraction of all Atlas members of the component belonging to any of the seven ESKAPE genera, and ESKAPE_genus_evenness is the normalised Shannon entropy of the distribution of Atlas members across those genera. S5 simultaneously rewards components that are both heavily enriched in AMR-genus proteins globally and concentrated in fewer genera rather than distributed evenly across all seven, distinguishing components with genuine AMR-clade association from those that happen to have moderate ESKAPE membership through broad taxonomic distribution. S5 is assigned a weight equal to S2 in Track B ($w_5$ = 0.25) because it captures the pathogen-concentration signal that species-level Atlas annotation alone cannot reliably provide for components spanning multiple AMR genera.

The DPPS for each component is thus computed as:

Track A: $DPPS = w_1S_1 + w_2S_2b + w_3S_3 + w_4S_4$    (weights: 0.15, 0.40, 0.25, 0.20)

$$\text{Track B: DPPS} = w_1S_1 + w_2S_2 + w_3S_3 + w_4S_4 + w_5S_5 \quad \text{(weights: 0.15, 0.25, 0.20, 0.15, 0.25)}$$

The robustness of component rankings to the assigned weight scheme was assessed through Monte Carlo sensitivity analysis (Section 2.6). Components were stratified into four priority tiers based on their DPPS: Tier I (DPPS ≥ 0.75), Tier II (0.50–0.75), Tier III (0.25–0.50), and Tier IV (< 0.25), with Tier I components designated as the highest-priority candidates for experimental follow-up.

**Algorithm 1. Pseudocode of ECLIPSE workflow**

```
Input:  query proteome Q (FASTA, all strains of target pathogen)
        Atlas AFDB90 network with precomputed community/component brightness
        strain count N
Output: dark components ranked by DPPS into priority tiers

# ---- Part I: Atlas mapping and darkness estimation ----
M ← MMseqs2_easy_search(Q, AFDB90, max_seqs=1)        # best Atlas hit per query
for each query Q in M:
    assign Q to its Atlas community and connected component
    inherit community brightness and component brightness
DARK ← { components with median component brightness = 0% }

# ---- taxonomic diversity of dark components ----
for each component c in DARK:
    ESKAPE_proportion(c)       ← fraction of Atlas members in AMR genera
    ESKAPE_genus_evenness(c)  ← normalised Shannon entropy across AMR genera
    ESKAPE_relative_evenness(c) ← normalised Shannon entropy (AMR vs non-AMR)

# ---- Part II: two-track stratification ----
TRACK_A ← { c in DARK : ESKAPE_proportion(c) = 1.0 and genus = target }   # pathogen-specific
TRACK_B ← { c in DARK : ESKAPE_proportion(c) ≥ 0.5 } \ TRACK_A            # ESKAPE-enriched

# ---- Part III: DPPS scoring ----
for each track T in {TRACK_A, TRACK_B}:
    T ← { c in T : median_length(c) ≥ 300 aa }           # length filter
    T ← MMseqs2_easy_cluster(T, id=0.30, cov=0.80)        # redundancy reduction
    for each component c in T:
        S1 ← 1 − brightness(c)/100                        # darkness
        S2 ← PA proportion in Atlas        (S2b = max(S2, strain_fraction) for Track A)
        S3 ← 1 − ESKAPE_relative_evenness(c)              # AMR-clade specificity
        S4 ← unique_strains(c) / N                        # strain coverage
        S5 ← ESKAPE_proportion(c) × (1 − ESKAPE_genus_evenness(c))   # Track B only
        DPPS(c) ← Σ wᵢ · Sᵢ                               # weighted sum, Σwᵢ = 1
    assign tiers: I (≥0.75), II (0.50–0.75), III (0.25–0.50), IV (<0.25)

# ---- robustness ----
for 500 Dirichlet-sampled weight vectors:
    recompute DPPS and record Tier I stability per component
```

return components ranked by DPPS with tier and stability

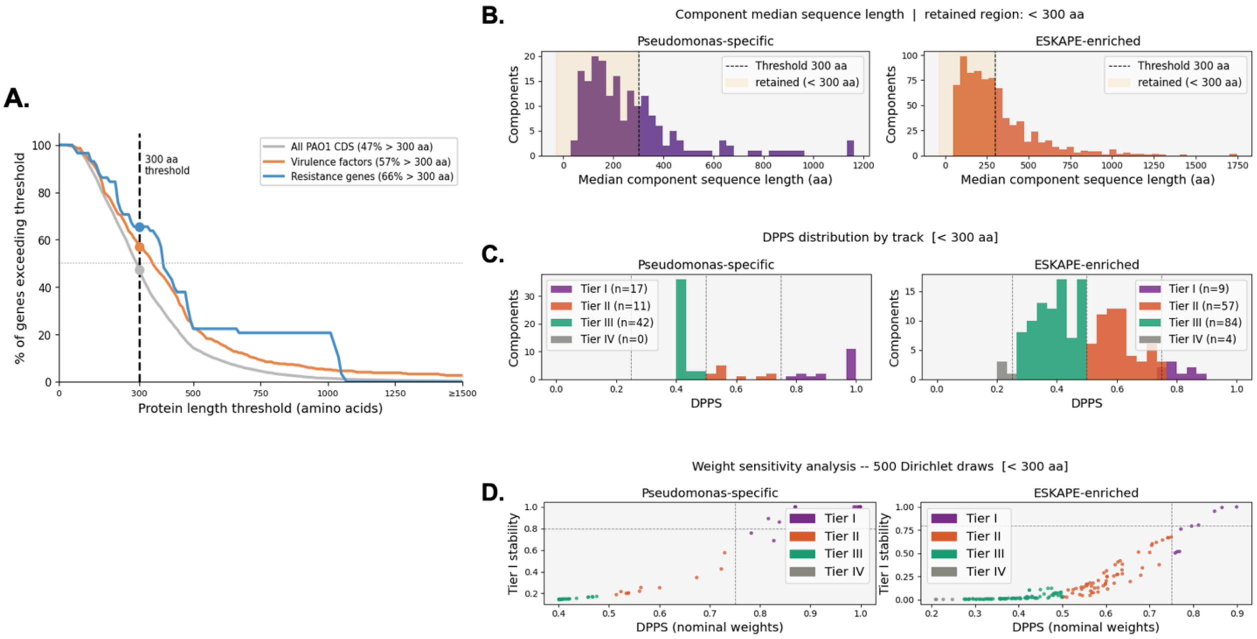


**Figure S1: A.** Length distribution of experimentally validated PAO1 virulence and resistance determinants relative to the 300 aa ECLIPSE length filter threshold. Cumulative distribution showing the percentage of each gene set exceeding the given length threshold; filled dots mark the exact percentage above 300 aa. **B.** Histograms showing the distribution of median sequence lengths below 300 aa across all dark components in the *Pseudomonas*-specific track (left, purple) and ESKAPE-enriched track (right, orange). **C.** Histograms show the distribution of composite DPPS values for all retained components in the *Pseudomonas*-specific track (left) and ESKAPE-enriched track (right), coloured by tier assignment: Tier **I** (purple, DPPS ≥ 0.75), Tier **II** (orange, 0.50–0.75), Tier **III** (teal, 0.25–0.50), and Tier IV (gray, < 0.25). **D.** Scatter plots show, for each component, the nominal DPPS score (x-axis) against its Tier **I** stability score (y-axis) the fraction of 500 Dirichlet-sampled random weight vectors under which the component achieved Tier **I** status (DPPS ≥ 0.75).

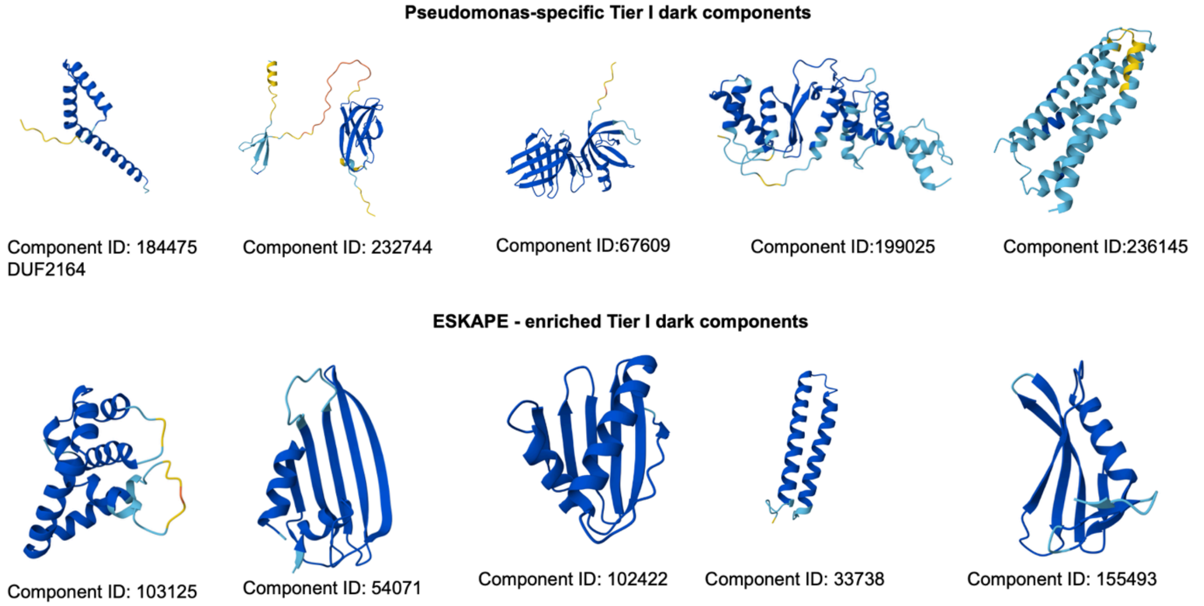


**Figure S2:** Alphafold2 modelled structure representatives for each Tier I candidate for *Pseudomonas*-specific and ESKAPE- enriched dark components of length below 300 aa respectively.

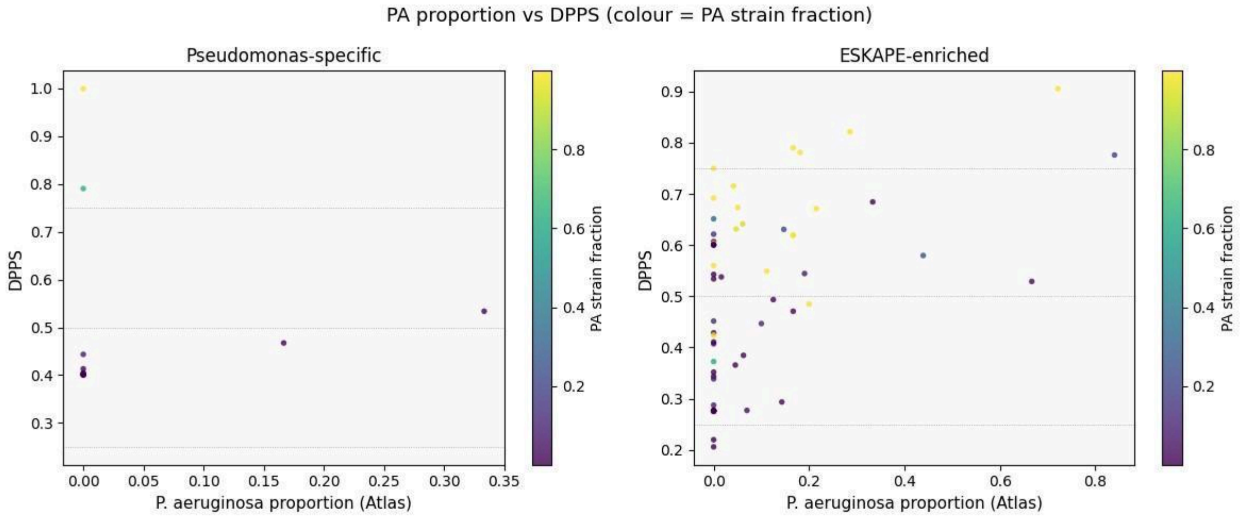


**Figure S3:** Scatter plots show the relationship between the *PA* proportion computed from the Atlas taxonomy files (x-axis) and the composite DPPS score (y-axis) for all components retained after the 300 aa median sequence length filter, for the *Pseudomonas*-specific track (left) and the ESKAPE-enriched track (right). Each point represents one dark connected component. Point colour indicates the *PA* strain fraction, the fraction of 635 query *PA* strains that contribute at least one protein to the component, computed directly from query protein identifiers, ranging from dark purple (low strain coverage) to bright yellow (near-universal conservation across the strain collection). Horizontal dotted lines mark the Tier I (DPPS = 0.75) and Tier II (DPPS = 0.50) tier boundaries. The vertical axis range differs between panels, reflecting the broader DPPS score range in the ESKAPE-enriched track.

In the *Pseudomonas*-specific track, many components cluster near zero Atlas *PA* proportion due to widespread non-canonical *Pseudomonas* sp. annotation in UniProt, yet several of these show high *PA* strain fraction (yellow points), confirming their genuine conservation across *PA* strains despite the annotation gap. In the ESKAPE-enriched track, the highest-scoring component (DPPS = 0.908, upper right, bright yellow) combines high Atlas *PA* proportion with near-universal strain coverage, representing the most robustly supported candidate across both independent measures of *PA* relevance. The divergence between Atlas *PA* proportion and *PA* strain fraction across both panels illustrates a fundamental limitation of species-level annotation in public databases and motivates the inclusion of the strain coverage sub-score S4 as an independent axis in the DPPS framework.

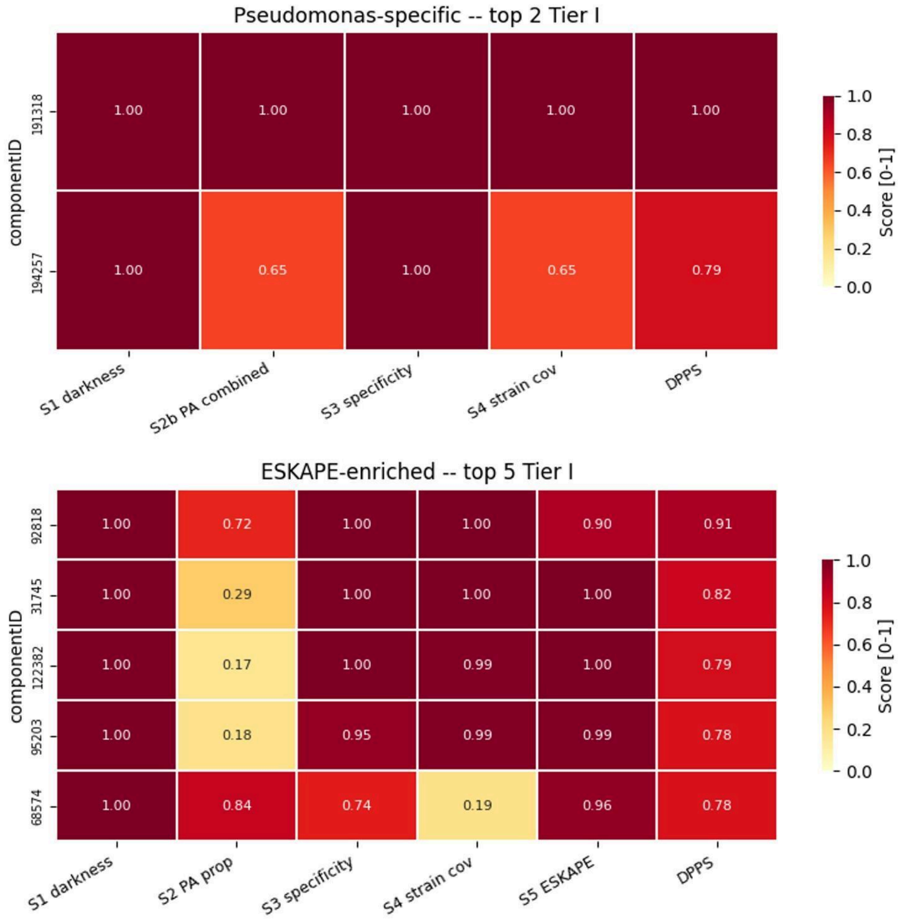


**Figure S4:** Heatmap showing the individual sub-score profile and composite DPPS for component 31745, the sole Tier I candidate in the *Pseudomonas*-specific track (DPPS = 0.76). Each column represents one scoring axis: S1 (darkness penalty), S2 (P. aeruginosa proportion from the Atlas), S3 (taxonomic specificity, defined as 1 − ESKAPE_relative_evenness), S4 (PA strain coverage, fraction of 635 query strains carrying the component), and the final composite DPPS. Cell values are annotated numerically. Colour intensity follows the scale from yellow (score = 0) to dark red (score = 1.0).Component 31745 achieves perfect scores on three of four sub-scores S1 = 1.00 (complete darkness across the Atlas), S3 = 1.00 (perfect AMR-clade restriction with no representation outside ESKAPE genera globally), and S4 = 1.00 (present in all 635 *P. aeruginosa* strains in the query collection). The only sub-score below maximum is S2 = 0.40, reflecting that 40% of Atlas UniRef50 members of this component are annotated as *PA* specifically, with the remaining members labelled as *Pseudomonas* sp. due to non-canonical species annotation in UniProt rather than reflecting genuine absence from PA. The composite DPPS of 0.76 exceeds the Tier I threshold of 0.75, driven by the convergence of universal strain conservation, perfect AMR-clade specificity, and complete functional darkness making this component a high-confidence candidate for experimental characterisation as a broadly conserved, pathogen-restricted, and completely uncharacterised *Pseudomonas*-specific protein family.

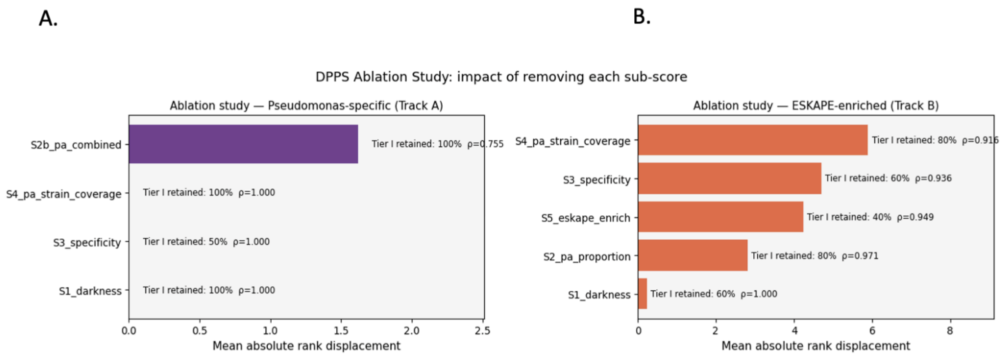


**Figure S5:** DPPS ablation analysis: Each bar shows the effect of removing one subscore for both track A and track B. Tier I candidates are retained in all the sub-scores with high Spearman rank correlation. In Track A, S2b (combined *PA* evidence) has the largest influence on rankings, while S4, S3, and S1 show minimal rank displacement with perfect rank correlation (ρ = 1.000), indicating that the *Pseudomonas*-specific ranking is primarily driven by *PA* evidence. In Track B, S4 (strain coverage) causes the greatest rank displacement upon removal, followed by S3 (AMR-clade specificity) and S5 (ESKAPE enrichment), demonstrating that the ESKAPE-enriched ranking is collectively determined by multiple sub-scores rather than any single dominant axis.

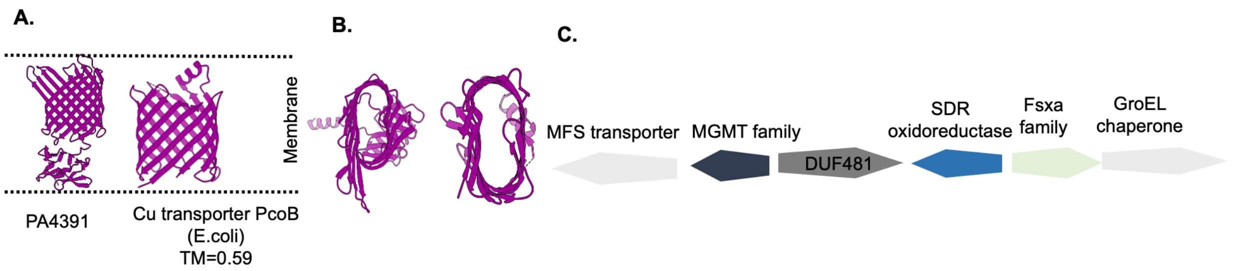


**Figure S6: A.** Predicted structure model of Tier II component 71592 (PA4391) with its closest foldseek hit Cu transporter from *PcoB*. **B.** Inner cross-section of component 71592 and *PcoB* foldseek hit. **C.** Genomic context analysis of flanking genes associated with component 71592.

| Sub-score | What it measures | Definition | Weight (Track A) | Weight (Track B) |
|---|---|---|---|---|
| **S1: Darkness** | Functional darkness of the component | 1 − (component brightness / 100); = 1.0 for all retained components | 0.15 | 0.15 |
| **S2b : Combined PA evidence** | PA-specificity within the *Pseudomonas* genus | Max (Atlas PA proportion, query strain fraction) | 0.40 | |
| **S2 : PA proportion** | PA representation from Atlas taxonomy | Fraction of Atlas UniRef50 members annotated as *P. aeruginosa* | | 0.25 |
| **S3: AMR-clade specificity** | Restriction to AMR genera globally | 1 − ESKAPE relative evenness | 0.25 | 0.20 |
| **S4: Strain coverage** | Empirical conservation across query strains | Fraction of 635 PA strains contributing ≥1 sequence | 0.20 | 0.15 |
| **S5 : ESKAPE enrichment** | Concentration within AMR genera | ESKAPE proportion × (1 − ESKAPE genus evenness) | | 0.25 |
| | | Total | 1.00 | 1.00 |

**Table S1: Summary of DPPS sub-scores, definitions, and track-specific weights.** Each component is scored by the Dark Proteome Prioritisation Score (DPPS), a weighted sum of normalised sub-scores. Four sub-scores are applied in the *Pseudomonas*-specific track (Track A) and five in the ESKAPE-enriched track (Track B). For each sub-score, the table lists the biological property it captures, its definition, and its weight in each track. All sub-scores are normalised to [0, 1] before weighting, and weights within each track sum to 1.0. The composite DPPS is computed as DPPS(Track A) = 0.15·S1 + 0.40·S2b + 0.25·S3 + 0.20·S4 and DPPS(Track B) = 0.15·S1 + 0.25·S2 + 0.20·S3 + 0.15·S4 + 0.25·S5. Components are stratified into four priority tiers based on DPPS: Tier I (≥ 0.75), Tier II (0.50–0.75), Tier III (0.25–0.50), and Tier IV (< 0.25), with Tier I designated the highest-priority candidates for experimental follow-up. Full sub-score derivations and biological rationale are provided in the Supplementary Methods.
